# The oocyte microenvironment is altered in adolescents compared to oocyte donors

**DOI:** 10.1101/2024.04.04.588118

**Authors:** Dilan Gokyer, Sophia Akinboro, Luhan T. Zhou, Anna Kleinhans, Monica M. Laronda, Francesca E. Duncan, Joan K. Riley, Kara N. Goldman, Elnur Babayev

**Affiliations:** Department of Obstetrics and Gynecology, Feinberg School of Medicine, Northwestern University, Chicago, IL, 60611; Weinberg College of Arts and Sciences, Northwestern University, Evanston, IL, 60208; Department of Obstetrics and Gynecology, Northwestern Medicine Center for Fertility and Reproductive Medicine, Chicago, IL, 60611; Department of Pediatrics, Feinberg School of Medicine, Northwestern University, Chicago, IL, 60611; Stanley Manne Children’s Research Institute, Ann & Robert H. Lurie Children’s Hospital of Chicago, Chicago, IL, 60611

**Keywords:** Cumulus cells, follicular fluid, fertility preservation, egg quality, children, RNA-seq, cytokine analysis

## Abstract

**Study question:** Are the molecular signatures of cumulus cells (CCs) and follicular fluid (FF) of adolescents undergoing fertility preservation differ from that of reproductively adult oocyte donors?

**Summary answer:** The microenvironment immediately surrounding the oocyte, including the CCs and FF, is altered in adolescents undergoing fertility preservation compared to oocyte donors.

**What is known already:** Adolescents experience a period of subfecundity following menarche. Recent evidence suggests that this may be at least partially due to increased oocyte aneuploidy. Reproductive juvenescence in mammals is associated with suboptimal oocyte quality.

**Study design, size, duration:** This was a prospective cohort study. Adolescents (10-19 years old, N=23) and oocyte donors (22-30 years old, N=31) undergoing ovarian stimulation and oocyte retrieval at the Northwestern Fertility and Reproductive Medicine Center between November 1, 2020 and May 1, 2023 were enrolled in this study.

**Participants/materials, setting, methods:** Patient demographics, ovarian stimulation, and oocyte retrieval outcomes were collected for all participants. The transcriptome of CCs associated with mature oocytes was compared between adolescents (10-19 years old, n=19), and oocyte donors (22-30 years old, n=19) using bulk RNA-sequencing. FF cytokine profiles (10-19 years old, n=18 vs. 25-30 years old, n=16) were compared using cytokine arrays.

**Main results and the role of chance:** RNA-seq analysis revealed 581 differentially expressed genes (DEGs) in cumulus cells of adolescents relative to oocyte donors, with 361 genes downregulated and 220 upregulated. Genes enriched in pathways involved in cell cycle and cell division (e.g., GO:1903047, p= 3.5 x 10^-43^; GO:0051983, p= 4.1 x 10^-30^; GO:0000281, p= 7.7 x 10^-15^; GO:0044839, p= 5.3 x 10^-13^) were significantly downregulated, while genes enriched in several pathways involved in cellular and vesicle organization (e.g., GO:0010256, p= 1.2 x 10^-8^; GO:0051129, p= 6.8 x 10^-7^; GO:0016050, p= 7.4 x 10^-7^; GO:0051640, p= 8.1 x 10^-7^) were upregulated in CCs of adolescents compared to oocyte donors. The levels of 9 cytokines were significantly increased in FF of adolescents compared to oocyte donors: IL-1 alpha (2-fold), IL-1 beta (1.7-fold), I-309 (2-fold), IL-15 (1.6-fold), TARC (1.9-fold), TPO (2.1-fold), IGFBP-4 (2-fold), IL-12-p40 (1.7-fold) and ENA-78 (1.4-fold). Interestingly, 7 of these cytokines have known pro-inflammatory roles. Importantly, neither the CC transcriptomes or FF cytokine profiles were different in adolescents with or without cancer.

**Large scale data:** Original high-throughput sequencing data will be deposited in Gene Expression Omnibus (GEO) before publication, and the GEO accession number will be provided here.

**Limitations, reasons for caution:** This study aims to gain insights into the associated gamete quality by studying the immediate oocyte microenvironment. The direct study of oocytes is more challenging due to sample scarcity, as they are cryopreserved for future use, but will provide a more accurate assessment of oocyte reproductive potential.

**Wider implications of the findings:** Understanding the underpinnings of altered immediate oocyte microenvironment of adolescent patients may provide insights into the reproductive potential of the associated gametes in the younger end of the age spectrum. This has implications for the fertility preservation cycles for very young patients.

**Study funding/competing interest(s):** This project was supported by Friends of Prentice organization SP0061324 (M.M.L and E.B.), Gesualdo Family Foundation (Research Scholar: M.M.L.), and NIH/NICHD K12 HD050121 (E.B.). The authors have declared that no conflict of interest exists.

## Introduction

Long-term survival of children with cancers has significantly improved in the last few decades due to advancements in oncology (Armstrong *et al*., 2016). However, many of the life-saving treatments are toxic to the gonads (Goldman *et al*., 2017). Patients with cancer report that fertility concerns cause significant distress to them and their family members, and many express a strong desire to preserve the possibility of having a biological child in future (Mulder *et al*., 2021). Similarly, patients with gender dysphoria prefer fertility preservation in some cases prior to gender affirming treatments due to unknown long-term effects of cross-sex hormone therapy on the quality of gametes or to prevent the detrimental psychological impacts of coming off of gender affirming hormones later in life (Chen and Simons, 2018; Moravek and Obedin-Maliver, 2021). Gamete, embryo, and ovarian/testis tissue cryopreservation are the available options to preserve fertility (ASRM Committee Opinion, 2019). Oocyte cryopreservation following controlled ovarian stimulation is the preferred method for fertility preservation for post-pubertal adolescents (ASRM Committee Opinion, 2021). Adolescence is defined as the phase of life between childhood and adulthood, from ages 10 to 19 (WHO, 2024).

Oocyte quantity and quality is highly dependent on age. With advanced reproductive age, there is a well-documented decrease in gamete quality due to oocyte aneuploidy and mitochondrial dysfunction as well as ovarian stromal inflammation and fibrosis (Wang *et al*., 2011; Duncan *et al*., 2012; Briley *et al*., 2016; Gruhn *et al*., 2019; Amargant *et al*., 2020; Beverley *et al*., 2021; Liu and Gao, 2023). Accumulating evidence also suggest that egg quality may be compromised at the other end of the age spectrum in very young individuals. The period of adolescent sterility or subfecundity is well documented in ethnology studies conducted in native island populations and in isolated communities in India and Thailand. In these communities, lower pregnancy rates were observed among young girls compared to adult women despite regular sexual intercourse in the absence of reliable contraception (Hartman, 1931; Ashley-Montagu, 1939; Wood and Milligan, 1989; Homan *et al*., 2007; Duncan, 2017). Similarly, a study of 42,493 parous, monogamously married, 19^th^ century women in the Utah Population Database reported that the natural fertility pattern in humans exhibits an inverse U-shaped curve where both young females (15 to early 20s) and women of advanced reproductive age (mid-30s and above) tend to experience lower fertility rates (Hawkes and Smith, 2010).

Decreased egg quality among very young mammals appears to be phylogenetically conserved (Duncan, 2017). Studies in mice, pig and non-human primate models demonstrate increased aneuploidy and/or decreased pregnancy rates in juvenescent animals (Mirskaia and Crew, 1931; Koenig and Stormshak, 1993; Wallen and Zehr, 2004; Lechniak *et al*., 2007; Kusuhara *et al*., 2021). Studies in humans also raise concerns of relatively poor gamete quality in children and young adults. A review of 15,169 trophectoderm biopsies demonstrates higher aneuploidy rates in patients in their early 20s (∼40% in women 22-23 year old) compared to the patients in their middle to late 20s (∼20-27% in women 26-30 years old) (Franasiak *et al*., 2014). Similarly, a recent study of oocytes from women undergoing ovarian tissue cryopreservation (9-39 years old) or oocytes and embryos from women undergoing IVF (20-43 years old) suggests a J-shaped aneuploidy curve in humans with increasing aneuploidy rates as age decreases below 27 years old (Gruhn *et al*., 2019).

Given the increasing number of fertility preservation cycles in children, adolescents, and young adults and concerns related to the suboptimal gamete quality in this age group, it is important to understand the quality of the cryopreserved gametes to counsel these patients on their reproductive potential, expected pregnancy outcomes, future offspring health, and design preventive and/or therapeutic strategies. Following ovarian stimulation, retrieved oocytes are vitrified for future use. However, the immediate microenvironment that surrounds the oocyte, which includes the cumulus cells (CCs) and follicular fluid (FF), can be readily sampled in IVF cycles. Importantly, these cells and biofluids reflect the quality of the associated gamete and key changes that occur with advanced reproductive age (Babayev and Duncan, 2022). CCs support the growth and development of the oocyte, are essential for fertility (Davis *et al*., 1999; Hizaki *et al*., 1999; Zhuo *et al*., 2001; Varani *et al*., 2002; Fülöp *et al*., 2003; Salustri *et al*., 2004), and undergo age-related genomic, transcriptomic, epigenomic, metabolomic, and proteomic changes (Lee *et al*., 2010; Tsai *et al*., 2010; McReynolds *et al*., 2012; Al-Edani *et al*., 2014; Molinari *et al*., 2016; Olsen *et al*., 2020; Babayev and Duncan, 2022). Similarly, FF reflects the metabolism, synthetic capacity and inflammatory signature of the surrounding granulosa and cumulus cells with advancing age (Adiga *et al*., 2002; Diez-Fraile *et al*., 2014; Machlin *et al*., 2021; Babayev and Duncan, 2022).

In this study, we tested the hypothesis that the biological profile of the oocyte immediate microenvironment changes as individuals transition from puberty to reproductive adulthood. To this end, we collected CCs, isolated from mature oocytes, and FF from adolescents undergoing fertility preservation and oocyte donors. We then compared the genome-wide CC transcriptomic signatures and the FF cytokine profile in these populations. To our knowledge, our adolescent group is the largest cohort to date with molecular analysis of the oocyte microenvironment at very young age. We chose oocyte donors as an older comparison group, because among the patients undergoing oocyte retrieval, they are a select group of presumably fertile adults with good ovarian reserve that represent a population with optimal egg quality. The observed alterations in the immediate oocyte microenvironment of adolescents, including dysregulated biological pathways in CCs and more proinflammatory cytokine signature in FF, may be reflective of the underlying differences in the gamete quality between these populations. These findings pave the way for our understanding of the reproductive potential of associated gametes in adolescents.

## Materials and Methods

### Population

Adolescent patients (10-19 years old, n=23) and oocyte donors (22-30 years old, n=31) undergoing ovarian stimulation and oocyte retrieval at the Northwestern Fertility and Reproductive Medicine Center between November 1, 2020 and May 1, 2023 were enrolled in this study. There were no exclusion criteria for these participants. Samples were collected from a single ovarian stimulation cycle for each participant. Age, race/ethnicity (as reported by the participant), past medical and surgical history, medication use, results of ovarian reserve and hormone laboratory testing, ovarian stimulation parameters, and the information on the number and maturation stage of oocytes was collected. All patients reported in this study had their demographics, medical history, IVF parameters and outcomes collected and analyzed. However, not all patients had CC and/or FF collected due to logistical reasons (e.g., embryology laboratory volume, staffing, research team availability). This IRB protocol continues to enroll patients to establish a biobank of cumulus cells and follicular fluid for future studies **(Supplementary Fig. S1)**.

### Ethical Approval

All participants gave written informed consent according to the protocol approved by Northwestern University Institutional Review Board (STU00213161).

### Sample Collection

N=19 adolescents and N=19 oocyte donors had high quality RNA extracted and sequencing libraries successfully prepared. Of all cumulus cell samples available at the time of RNA extraction, only one (adolescent) failed successful RNA extraction and library preparation. For FF cytokines arrays, we analyzed all adolescent FF samples available at the time these arrays were performed (n=18), however, we only included donors that were 25-30 years old (n=16) due to the setup of these arrays to maintain a relatively large age gap between adolescent and oocyte donor samples **(Supplementary Fig. S1)**.

CCs were mechanically dissected from cumulus-oocyte complexes (COCs) prior to hyaluronidase treatment to avoid changes in gene expression in the associated cells due to disruption of the extracellular matrix **(Figure 1C)** (Spencer *et al*., 2007a; Lelièvre, 2009; Hastings *et al*., 2019). Our method ensures that the CC transcriptome is preserved in its native state. CC masses from 6-12 COCs per participant were collected. Four CC masses dissected from each expanded COC were immediately washed in 500 μL Phosphate-buffered solution (PBS) (Cat #20012027, Fisher Scientific, Waltham, MO) and snap frozen in 20 μL of RNA later (Cat #AM7024, Fisher Scientific, Waltham, MO**) (Figure 1C, Supplementary Fig. S2A).** Microdissected COCs were rinsed and held for ∼2 hours in Quinn’s Advantage Fertilization Medium supplemented with 5% human serum albumin (Cat# ART-1021 and ART-3001, CooperSurgical Fertility Companies, Denmark) in a 37°C, 6% CO2, and 5% O2 incubator until denudation as per routine IVF laboratory protocol **(Figure 1C, Supplementary Fig. S2B).** Each corresponding COC was tracked during CC stripping. After denudation, the maturation stage of the oocyte was determined based on morphology and linked to the collected CC samples. Oocytes arrested at metaphase of meiosis II (MII) were characterized by extrusion of the first polar body (PBI), whereas oocytes that failed to mature and remained arrested at prophase of meiosis I were characterized by an intact nucleus or germinal vesicle (GV oocyte). Oocytes that had undergone germinal vesicle breakdown but lacked PBI (in between prophase I and MII) were referred to as a MI oocyte. Since the maturity of the oocyte influences CC gene expression (Luciano *et al*., 2000), we only used CCs from mature (MII) oocytes for further analysis. All CC samples were stored at -80°C until further processing for RNA-seq.

**Figure 1.**
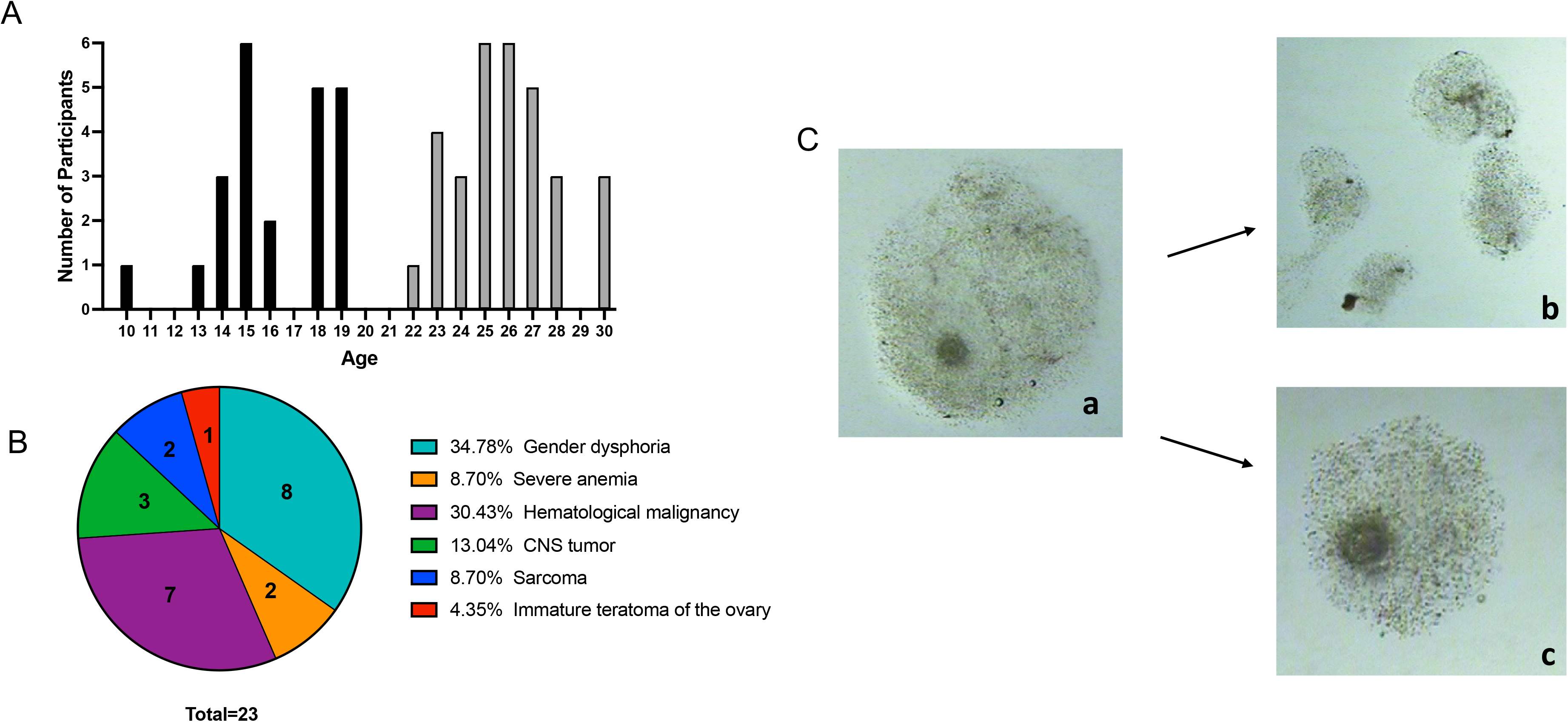
Participant age and medical diagnosis, and cumulus cell collection. A) Distribution of age among adolescents (black bars) and oocyte donors (grey bars) B) Medical diagnoses of adolescents C) Microdissection of cumulus cells from retrieved expanded cumulus-oocyte complexes (COCs) prior to hyaluronidase treatment: a - expanded COC before microdissection of cumulus cell masses, b - trimmed cumulus cell masses (n=4), c - COC after microdissection.

Follicular fluid samples from the two dominant follicles (18-20 mm) of each ovary was collected per participant. Halt protease inhibitor cocktail (Cat #87786, Thermo Fisher Scientific) was added at 1X concentration to avoid protein degradation. Cells were pelleted via centrifugation at 380 × g for 10 min at 4 °C, and the FF supernatant was transferred to sterile 1.5 ml Eppendorf tubes, and samples were stored at -80^°^C until further analysis.

### RNA Isolation and Sequencing

Our initial experiments demonstrated that the combination of CCs from 3 MII oocytes provides the optimal yield and concentration of RNA. Therefore, CC samples from 3 MII oocytes per participant were combined, and RNA was isolated for all study participants in parallel using an RNAeasy Micro Kit (Cat #74004, Qiagen, Valencia, CA). Extracted RNA was quantified with a Qubit fluorometer (Cat #Q33238, Thermo Fisher Scientific), and RNA quality control was performed with a Bioanalyzer RNA 6000 Pico Chip (Cat #5067-1513, Agilent Technologies, CA). Samples with an RNA Integrity Number (RIN) >7 passed quality control and were used for library preparation with the Illumina Stranded mRNA Library Prep Kit according to manufacturer’s instructions (Cat #20040532, Illumina Inc., San Diego, CA). This procedure includes mRNA purification and fragmentation, cDNA synthesis, 3’ end adenylation, Illumina adapter ligation, library PCR amplification and validation. The lllumina NovaSeq 6000 sequencer (Cat #20068232, Illumina Inc., San Diego, CA) was used to sequence the libraries with the production of paired-end 150 base 20-25 million reads per sample on an S4 flow cell.

### Bioinformatic Analysis

The quality of reads, in FASTQ format, was evaluated using FastQC. Reads were trimmed to remove Illumina adapters from the 3’ ends using cutadapt (Spencer *et al*., 2007b). Trimmed reads were aligned to the Homo sapiens genome (hg38) using STAR (Dobin *et al*., 2013). Read counts for each gene were calculated using htseq-count (Anders *et al*., 2015), and in conjunction with a gene annotation file consisted of 58396 genes for hg38 obtained from Ensembl (Martin *et al*., 2023). Normalization and differential expression were calculated using DESeq2 which employs the Wald test (Love *et al*., 2014). The cutoff for determining significantly differentially expressed genes (DEGs) was an FDR-adjusted p-value less than 0.05 using the Benjamini-Hochberg method. Enrichment analysis was performed using downregulated and upregulated DEGs via Metascape online platform (Zhou *et al*., 2019) to identify differences in gene ontology (GO) biological pathways across two groups. Top 20 significantly enriched GO terms were plotted using RStudio version 4.3.1 (R Foundation for Statistical Computing, Vienna, Austria, https://www.R-project.org/).

### Cytokine Antibody Array

FF samples from adolescent and oocyte donors were thawed and a 2-fold dilution was performed using the blocking buffer provided in the Human Cytokine Array C5 as previously described (Ray Biotech Inc, Norcross, GA, USA) (Machlin *et al*., 2021). 1mL of diluted samples were run as duplicates in parallel on arrays according to the manufacturer’s instructions. The arrays were visualized by chemiluminescence using a BioRad ChemiDoc Imaging System (Cat #12003153, Bio-Rad Laboratories Inc., Hercules, CA). The resulting chemiluminescence data were quantified using the Protein Array Analyzer plugin for FIJI software (Schindelin *et al*., 2012). The relative intensity units (RU) of 80 cytokines were averaged between duplicate FF samples with the background subtracted. The RU of these 80 cytokines was compared between adolescents and oocyte donors.

### Statistical Analysis

The normal distribution of the data was evaluated with the Shapiro-Wilk and Kolmogorov-Smirnov test. Analysis between two groups of continuous variables were performed with unpaired two-sided Student’s t-test or Mann-Whitney U test depending on distribution. Categorical variables were analyzed with Fisher’s exact test or Chi-square test. Data are presented as mean ± SEM. P values < 0.05 were considered statistically significant. GraphPad Prism version 9.0.1 (Boston, Massachusetts USA, www.graphpad.com) was used for statistical analysis.

## Results

### Demographic and clinical characteristics of participants

Adolescent participants were on average a decade younger than oocyte donors (16.6 ± 0.5, 10-19 years old, n=23 vs. 26.3 ± 0.4, 22-30 years old, n=31, p<0.0001) **(Table 1**, **Figure 1A)**. Thirteen adolescents were diagnosed with cancer prior to fertility preservation, whereas ten adolescents had a non-cancer diagnosis, including gender dysphoria (n=8) and severe anemia requiring stem cell therapy (n=2) **(Figure 1B).** Five of thirteen patients received previous chemo- and/or radiotherapy. BMI (25.2 ± 1.35 vs. 24.2 ± 0.45 kg/m^2^, p=0.83) and race/ethnicity were similar across groups. Only one adolescent patient (10 years old) was pre-menarchal, whereas the rest (n=22) were at least 2 years post-menarchal. As expected, Anti-Mullerian hormone (AMH) levels and antral follicle count (AFC) were significantly higher in oocyte donors compared to adolescents as donors represent a select group of women with good ovarian reserve (AMH: 6.6 ± 0.66 ng/ml vs. 3.3 ± 0.47 and AFC: 26.3 ± 1.8 vs. 16.8 ± 1.2, p<0.0001) **(Table 1).** In line with this, oocyte donors required lower total gonadotropin doses (4131 ± 274 vs. 5322 ± 427, p=0.024) and had higher peak estradiol (E2) levels (3820 ± 256 vs. 2450 ± 256, p=0.0006) compared to adolescents. Luteal phase start (34.78% vs. 3.23%, p=0.003) and abdominal ultrasound monitoring (65% vs 0%, P<0.0001) were more common for adolescents. Duration of stimulation (11.3 ± 0.3 vs. 11.3 ± 0.2, p=0.92), the number of monitoring visits (6.1 ± 0.2 vs. 6.6 ± 0.2, p=0.14), and the number of retrieved oocytes (MII: 20.2 ± 2.6 vs. 24.4 ± 2.3, p=0.15; MI: 1.3 ± 0.3 vs. 2.2 ± 0.4, p=0.09 and GV: 4.5 ± 1.1 vs. 2.8 ± 0.6, p=0.26) were similar between groups. The number of degenerate gametes (1.6 ± 0.4 vs. 0.5 ± 0.2, p<0.05) as well as empty zonae collected (EZ) (2.0 ± 0.6 vs. 0.4 ± 0.2, p<0.01) at retrieval were higher in adolescent participants, but this finding is likely clinically insignificant given the low absolute numbers.

**Table 1.** Demographics and IVF cycle characteristics.

|  | **Adolescents**<br>**(n=23)** | **Donors**<br>**(n=31)** | **P value** |
| --- | --- | --- | --- |
| **Age (years)** | 16.6 ± 0.5 | 26.3 ± 0.4 | <0.0001 |
| **BMI (kg/m<sup>2</sup>)** | 25.2 ± 1.35 | 24.2 ± 0.45 | 0.8317 |
| **Race/ethnicity** (number of participants) |  |  | 0.3717 |
| Caucasian | 12 | 25 |  |
| African American | 3 | 2 |  |
| Asian | 1 | 2 |  |
| Middle Eastern | 1 | 0 |  |
| Hispanic | 1 | 1 |  |
| Multi-racial | 3 | 1 |  |
| **AMH (ng/mL)** | 3.3 ± 0.47 | 6.6 ± 0.66 | <0.0001 |
| **Antral follicle count (AFC)** | 16.8 ± 1.2 | 26.3 ± 1.8 | <0.0001 |
| **Luteal phase start** | 34.78% | 3.23% | 0.003 |
| **Duration of stimulation (days)** | 11.3 ± 0.3 | 11.3 ± 0.2 | 0.9248 |
| **Number of monitoring visits (days)** | 6.1 ± 0.2 | 6.6 ± 0.2 | 0.1382 |
| **Type of ultrasound** |  |  | <0.0001 |
| Transvaginal% | 34.78% | 100.00% |  |
| Transabdominal% | 65.22% |  |  |
| **Total gonadotropin dose (IU)** | 5322 ± 427 | 4131 ± 274 | 0.024 |
| **Peak estradiol (pg/mL)** | 2450 ± 256 | 3820 ± 256 | 0.0006 |
| **Number of oocytes** | 29.9 ± 3.8 | 32.7 ± 2.3 | 0.5088 |
| **Number of MII**s | 20.2 ± 2.6 | 24.4 ± 2.3 | 0.1461 |
| **Number of MI**s | 1.3 ± 0.3 | 2.2 ± 0.4 | 0.0896 |
| **Number of GV**s | 4.5 ± 1.1 | 2.8 ± 0.6 | 0.2606 |
| **Number of degenerated oocytes at retrieval** | 1.6 ± 0.4 | 0.5 ± 0.2 | 0.0195 |
| **Number of EZ**s | 2.0 ± 0.6 | 0.4 ± 0.2 | 0.0007 |

Values are presented as mean ± SEM. BMI = body mass index; AMH = Anti-Mullerian hormone; MII = mature metaphase II arrested oocytes; MI = immature oocytes between GV and MII stage; GV = immature oocytes with germinal vesicle; EZs = zona pellucida devoid of an oocyte.

### The cumulus cell transcriptome of adolescents reveals differentially regulated biological pathways compared to oocyte donors

Participants included in the CC transcriptome analysis had similar demographic and IVF cycle parameters to the overall cohort **(Supplementary Fig. S3)** with average 8 year age gap between groups (16.4 ± 0.5 years old, n=19 vs. 24.1 ± 0.6 years old, n=19). Unsupervised hierarchical clustering and Principal component analysis (PCA) did not demonstrate clear clustering of cumulus cell transcriptomes from adolescents and oocyte donors indicating overall similar cumulus cell biology in these age groups which is not surprising giving highly differentiated nature of these cells **(Supplementary Fig. S4).** However, comparative transcriptomic analysis revealed 581 DEGs **(Figure 2A).** 361 genes were downregulated and 220 were upregulated in cumulus cells of adolescents compared to oocyte donors **(Figure 2A, B)**. Genes enriched in biological pathways involved in cell cycle and cell division processes were significantly downregulated in adolescents compared to oocyte donors. Mitotic cell cycle process (GO:0051983, p= 4.1 x 10^-30^), regulation of chromosome segregation (GO:0051983, p= 3.5 x 10^-43^), positive regulation of cell cycle process (GO:0090068, p= 4 x 10^-30^), mitotic cytokinesis (GO:0000281, p= 7.7 x 10^-15^), cell cycle G2/M phase transition (GO:0044839, p= 5.2 x 10^-13^), and chromosome condensation (GO:0030261, p= 3.6 x 10^-10^) were top down-regulated biological pathways **(Figure 2C).** Transcripts in pathways involved in cellular communication, intracellular transport processes and organelle organization were more abundant in CCs of adolescents compared to oocyte donors. Endomembrane system organization (GO:0010256, p= 1.2 x 10^-8^), sterol biosynthetic process (GO:0016126, p= 5.6 x 10^-7^), negative regulation of cellular component organization (GO:0051129, p= 6.8 x 10^-7^), vesicle organization (GO:0016050, p= 7.4 x 10^-7^), organelle organization (GO:0006886, p= 8 x 10^-7^), and intracellular protein transport (GO:0016050, p= 1.9 x 10^-6^) were top up-regulated biological pathways **(Figure 2D).**

**Figure 2.**
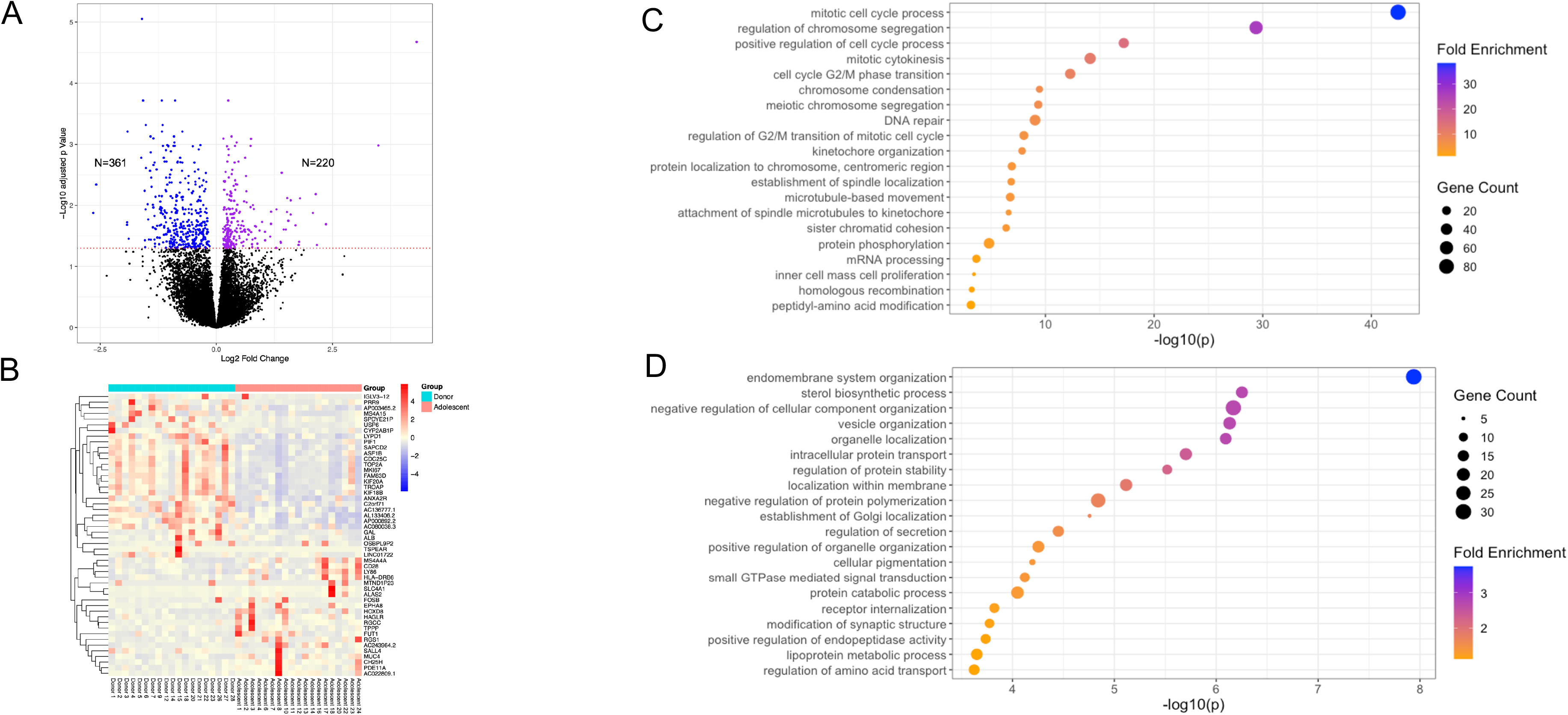
Comparative RNA-seq analysis of cumulus cells collected from adolescents and oocyte donors. A) Volcano plot with downregulated (n=361) and upregulated DEGs (n=220) in adolescents (n=19) compared to donors (n=19) (dashed red line - adjusted p<0.05). B) Heatmap with the top 50 DEGs with the highest fold change between two age groups. C) Top 20 significantly downregulated GO terms (biological pathways) in adolescents compared to donors. D) Top 20 significantly upregulated GO terms (biological pathways) in adolescents compared to donors. DEGs: differentially expressed genes.

Several subgroup analyses were performed to ascertain the impact of age and cancer diagnosis on DEGs and altered biological pathways. Although the number of DEGs between CCs from adolescents and oocyte donors increased to 1069 when the analysis was limited to individuals <16 years old (median age of participants), the biological pathways affected overall remained similar to the original analysis (**Supplementary Fig. S5).** Furthermore, CC transcriptomic analysis of adolescents ≥16 years old (n=10) vs. <16 (n=9) revealed only 41 DEGs **(Supplementary Fig. S6).** These results demonstrate that biological pathways appear dysregulated in cumulus cells of adolescents of all ages compared to oocyte donors in our study. Importantly, no DEGs were identified in cumulus cells of adolescents with cancer (n=11) compared to adolescents without cancer (n=8) **(Supplementary Fig. S7)** indicating that the observed molecular differences are most likely attributable to age and not the underlying cancer diagnosis.

### Follicular fluid of adolescents is more pro-inflammatory compared to oocyte donors

FF cytokine analysis of adolescents (16.7 ± 0.6, 10-19 years old, n=18) and oocyte donors (27.3 ± 0.4, 25-30 years old, n=16) **(Supplementary Fig. S8)** revealed altered levels of 9 cytokines **(Figure 3A).** All 9 cytokines demonstrated higher levels in FF of adolescents compared to oocyte donors: Interleukin-1 alpha (IL-1α; 2-fold), Interleukin-1 beta (IL-1β; 1.7-fold), I-309 (2-fold), Interleukin-15 (IL-15; 1.6-fold), Thymus and activation-regulated chemokine (TARC; 1.9-fold), thrombopoietin (TPO; 2.1-fold), Insulin-like growth factor binding protein-4 (IGFBP-4; 2-fold), the p40 Subunit of Interleukin-12 (IL-12-p40; 1.7-fold) and Epithelial neutrophil-activating protein 78 (ENA-78; 1.4-fold) **(Figure 3B)**. Interestingly, 7 of these 9 cytokines have pro-inflammatory roles (Miller and Krangel, 1992; Trinchieri, 1995; Gilchrest *et al*., 2003; Persson *et al*., 2003; Perera *et al*., 2012; Santana Carrero *et al*., 2019; Galozzi *et al*., 2021; Lupancu *et al*., 2023) suggesting a more pro-inflammatory environment in the FF of the adolescents compared to oocyte donors. The other two, IGFBP-4 and TPO, were shown to be important in the regulation of insulin-like growth factor (IGF) action (Laursen *et al*., 2007) and in the production of platelets (Hitchcock *et al*., 2021), respectively. None of these 9 cytokines demonstrated altered levels in FF of adolescents with cancer (n=12) compared to those without cancer diagnosis (n=6) **(Supplementary Fig. S9).** FF of adolescent patients with cancer had decreased levels of 9 cytokines (G-CSF, EGF, FGF-7, BDNF, BLC, LIGHT, MCP-2, IL-7, TGF-beta 1) and increased levels of one cytokine (MCP-1) compared to FF of adolescents without cancer **(Supplementary Fig. S9)**. This demonstrates that increased levels of pro-inflammatory cytokines in FF of adolescents is likely due to younger age and not the cancer diagnosis.

**Figure 3.**
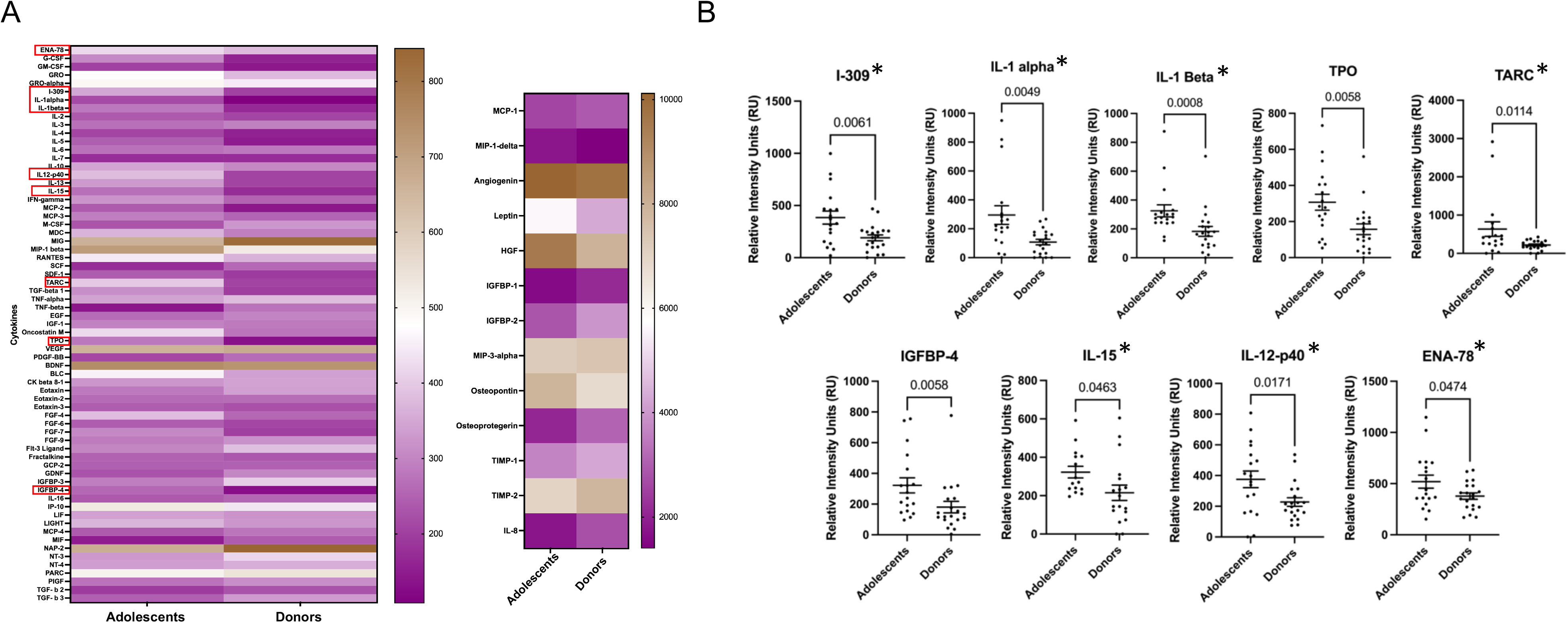
Follicular fluid levels of cytokines in adolescents and oocyte donors. A) Heatmap of follicular fluid levels of 80 cytokines in adolescents and oocyte donors with 9 of them significantly different between the two groups (red boxes) B) 7 of these 9 significantly different cytokines have pro-inflammatory roles (asterisks).

## Discussion

Oocyte cryopreservation is an essential fertility preservation option for adolescents and young adults. However, the quality and reproductive potential of the gametes in this age group is understudied despite concerns of potential suboptimal gamete quality (Franasiak *et al*., 2014; Gruhn *et al*., 2019). Moreover, the outcomes of pregnancies with these oocytes as well as the health of future generations is unknown. Our unbiased molecular analysis of the immediate oocyte microenvironment, in the largest cohort of adolescents to date, revealed dysregulated biological pathways in CCs and a more pro-inflammatory signature in FF compared to oocyte donors. These alterations in the immediate oocyte microenvironment likely reflect the biology of the underlying oocyte and provide potential targets for modulation to improve reproductive outcomes of adolescent fertility preservation cycles.

Oocytes are dependent on the surrounding cumulus cells (Gilchrist *et al*., 2004). CCs are interconnected with oocytes through gap junctions located at transzonal projections, forming a cohesive cumulus-oocyte complex (COC) (Xie *et al*., 2023). Oocytes exert control over CCs’ DNA synthesis, proliferation, differentiation, expansion, and prevent their apoptosis with oocyte-secreted factors (OSFs) (Hussein *et al*., 2005; Gilchrist *et al*., 2008; Cakmak *et al*., 2016; Richani *et al*., 2022). The regulation of cumulus cell function in response to these signals affects the oocyte’s competency via bidirectional communication to support the subsequent fertilization and embryonic development (Gilchrist *et al*., 2008). In preovulatory follicles, luteinizing hormone (LH) surge triggers synthesis and secretion of epidermal growth factor (EGF)-like mediators from mural granulosa cells (MGCs). As oocyte and CCs lack LH receptors, these factors including amphiregulin (AREG), epiregulin, and beta-cellulin propagate the LH stimulus down to COCs (Park *et al*., 2004). These mediators augment cumulus cell proliferation in mice, goat, and pigs (Cakmak *et al*., 2016; Baszary and Moniharapon, 2020; Zhang *et al*., 2022). A study in a bovine model demonstrated that the interruption of the communication axis between oocyte and cumulus cells by dissociation of CCs from COCs results in decreased cumulus cell proliferation and expansion, ultimately triggering apoptosis in both cell types (Luciano *et al*., 2000). Down-regulated expression of proliferation pathways in adolescent cumulus cells may indicate abnormal response to or disordered signaling of EGF-like growth factors, and the upregulation of pathways related to cellular organization, transport, and intercellular communication may signify a compensatory mechanism. In fact, some of the genes involved in EGF signaling demonstrate altered transcript levels in cumulus cells of adolescents compared to oocyte donors (i.e., *TMEFF1, ZGPAT, MAPK8IP3, PBK*) which warrants further investigation. Whether these molecular differences lead to impaired cumulus expansion, similar to what is observed with advanced reproductive age (Babayev *et al*., 2023), also remains to be investigated.

It is plausible that the differences in the phase of menstrual cycle at the start of hormonal stimulation, gonadotropin doses, and peak estradiol levels – donors not unexpectedly required less gonadotropin stimulation and had higher estradiol levels at the time of trigger – may have contributed to the observed molecular disparities between our groups. However, other demographic (e.g., BMI, race/ethnicity) and oocyte retrieval parameters (e.g., trigger criteria, number of oocytes at different stages of maturation) were similar between the groups. Furthermore, our study design ensured that the cumulus cell samples were collected only from COCs with mature oocytes retrieved from large antral follicles, and follicular fluid samples were collected from large antral follicles in both groups. Therefore, age was likely the primary variable driving the observed alterations in the immediate oocyte microenvironment.

Importantly, no DEGs were noted between adolescents with cancer compared to those without. This suggests that the observed molecular differences are likely attributed to age rather than the presence of cancer. We also cannot exclude that the previous chemo- and/or radiotherapy (5/13 adolescent patients) may have affected our results. However, given no DEGs in cumulus cells of adolescents with versus without cancer, this is unlikely to be a major contributor to the dyregulated gene expression observed in the adolescent group compared to the oocyte donors. Interestingly, the affected biological pathways remained generally similar when the analysis was limited to adolescents <16 years old compared to oocyte donors, and very few DEGs were observed between adolescents ≥16 years old compared to those <16 years old. This suggests that oocyte cryopreservation in an adolescent population may have similar prognosis across an age spectrum of 10 to 19 years old. However, larger studies directly examining the quality of oocytes are needed for definitive conclusions. Another limitation of our study is the reliance on the analysis of cumulus cell transcriptome, and future studies focusing on the proteome of these cells will elucidate whether observed dysregulation persists at the protein level.

Follicular fluid reflects the metabolism of the surrounding granulosa and cumulus cells and plays a crucial role in supporting the maturation and development of the oocyte within the antral follicle (Edwards, 1974). Reproductive aging is associated with inflammation, oxidative damage, ovarian fibrosis, and increased stiffness in the ovarian stroma (Amargant *et al*., 2020; Wang *et al*., 2021; Umehara *et al*., 2022; Babayev *et al*., 2023). Similarly, follicular fluid demonstrates a more fibroinflammatory cytokine signature (IL-3, IL-7, IL-15, TGFβ1, TGFβ3, and MIP-1) with aging (Machlin *et al*., 2021). Our study demonstrated that follicular fluid of adolescents has higher levels of proinflammatory cytokines (IL-1α, IL-1β, I-309, IL-15, TARC, IL-12-p40, ENA-78) compared to oocyte donors. Majority of these cytokines were unique to adolescent population and notably, only one of these cytokines, IL-15, was shared with women of advanced reproductive age reported in the previous study (Machlin *et al*., 2021). Although these data may indicate that the tendency for more proinflammatory milieu may exist at both ends of the age spectrum, the levels of individuals cytokines and mechanisms may differ. Cancer is associated with inflammation (Singh *et al*., 2019). However, none of the aforementioned cytokines had significantly different levels in FF of adolescents with cancer compared to those without the cancer diagnosis. This indicates that the increase in proinflammatory cytokine levels in FF of adolescents compared to donors is likely attributed to age rather than cancer diagnosis.

The success of the bidirectional regulatory loop between oocytes and CCs, and the oocyte’s intrinsic ability to govern its own microenvironment by OSFs are crucial characteristics contributing to the oocyte quality (Corn *et al*., 2005). Therefore, more proinflammatory follicular fluid environment and dysregulation in cumulus cell proliferation and organelle organization pathways could possibly be attributed to the suboptimal oocyte quality in adolescents. Our study provides insights into the molecular processes associated with the altered oocyte microenvironment in adolescents, and paves the way for further research to deepen our understanding of the reproductive potential of cryopreserved oocytes in this age group.

### Authors’ roles

E.B., K.N.G., M.M.L. and F.E.D. conceived the original idea. E.B. designed the experiments. D.G., S.A., L.T.Z, A.K., J.K.R., E.B. carried out the experiments. D.G., K.N.G., M.M.L., F.E.D., J.K.R and E.B. analyzed and interpreted the data. D.G. and E.B. wrote the manuscript. M.M.L., F.E.D., J.K.R. and K.N.G. provided critical discussion, reviewed, and revised the manuscript.

## Acknowledgements

This work was supported by the Northwestern University NUSeq Core Facility.

