## Supplementary Figure for "The oocyte microenvironment is altered in adolescents compared to oocyte donors"

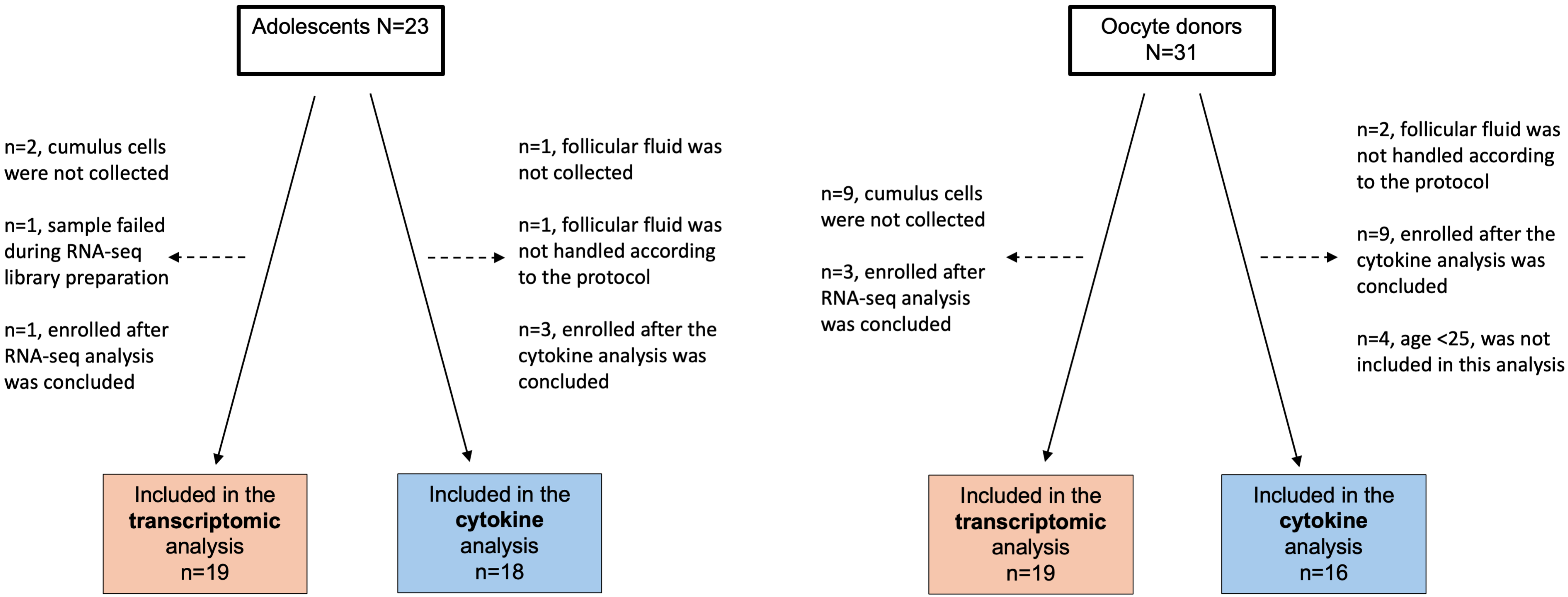


**Supplementary Figure 1. Study population and sample collection scheme**

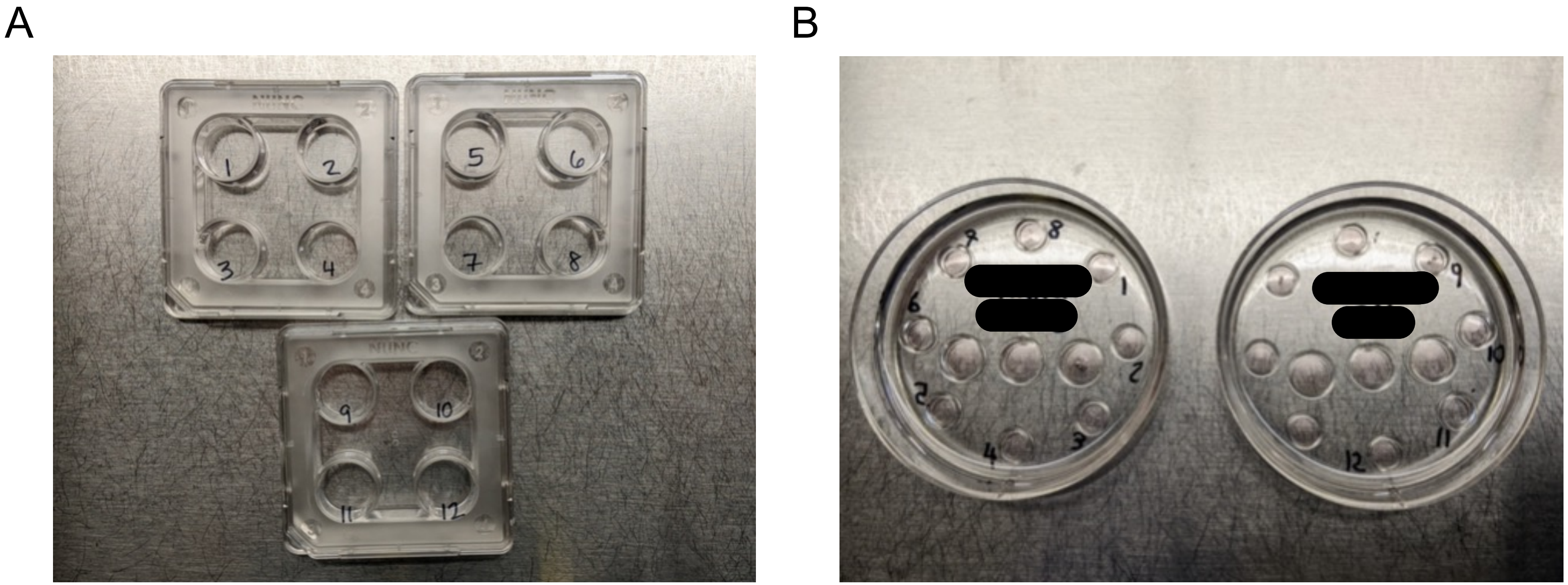


**Supplementary Figure 2. Plate set-ups used after microdissection of cumulus cells from expanded cumulus-oocyte complexes (COCs) prior to hyaluronidase treatment.** A) 4-well plate setup: cumulus cell masses are rinsed in PBS B) Plate setup: microdissected COCs are rinsed in Quinn's Advantage Fertilization Medium supplemented with 5% human serum albumin and held in the incubator for ~2 hours until denudation.

A

| | Adolescents (n=19) | Donors (n=19) | P value |
|---|---|---|---|
| **Age** (years) | 16.4 ± 0.5 | 24.1 ± 0.6 | <0.0001 |
| **BMI** (kg/m$^2$) | 25.8 ± 1.54 | 24.1 ± 0.56 | 0.598 |
| **Race/ethnicity** (number of participants) | | | 0.4662 |
| Caucasian | 9 | 15 | |
| African American | 3 | 2 | |
| Asian | 1 | 1 | |
| Middle Eastern | 1 | 0 | |
| Hispanic | 1 | 1 | |
| Multi-racial | 2 | 0 | |
| **AMH** (ng/mL) | 3.4 ± 0.56 | 6.9 ± 0.89 | 0.0009 |
| **Antral follicle count (AFC)** | 16.1 ± 1.2 | 27.8 ± 2.8 | <0.0001 |
| **Luteal start** | 42.11% | 5.26% | 0.0188 |
| **Duration of stimulation** (days) | 11.3 ± 0.3 | 11.5 ± 0.2 | 0.3663 |
| **Number of monitoring visits** (days) | 6.2 ± 0.2 | 6.8 ± 0.3 | 0.099 |
| **Type of USG** | | | <0.0001 |
| Transvaginal% | 31.58% | 100.00% | |
| Transabdominal% | 68.42% | | |
| **Total gonadotropin dose** (IU) | 5341 ± 442 | 3968 ± 303 | 0.0171 |
| **Peak estradiol** (pg/mL) | 2422 ± 303 | 4034 ± 356 | 0.0009 |
| **Number of oocytes** | 31 ± 4.1 | 31.8 ± 2.7 | 0.8733 |
| **Number of MIIs** | 21 ± 2.8 | 25.8 ± 2.6 | 0.124 |
| **Number of MIs** | 1.5 ± 0.3 | 2.6 ± 0.4 | 0.0274 |
| **Number of GVs** | 4.1 ± 1.2 | 2.8 ± 0.6 | 0.7095 |
| **Number of degenerated oocytes at retrieval** | 1.7 ± 0.5 | 0.2 ± 0.1 | 0.0058 |
| **Number of EZs** | 2.3 ± 0.7 | 0.4 ± 0.2 | 0.0037 |

Values are presented as mean ± SEM. BMI = body mass index; AMH = Anti-Mullerian hormone; MIIs = mature metaphase II arrested oocytes; MIs = immature oocytes between GV and MII stage. GVs = immature oocytes with germinal vesicle; EZs = zona pellucida devoid of an oocyte.

B

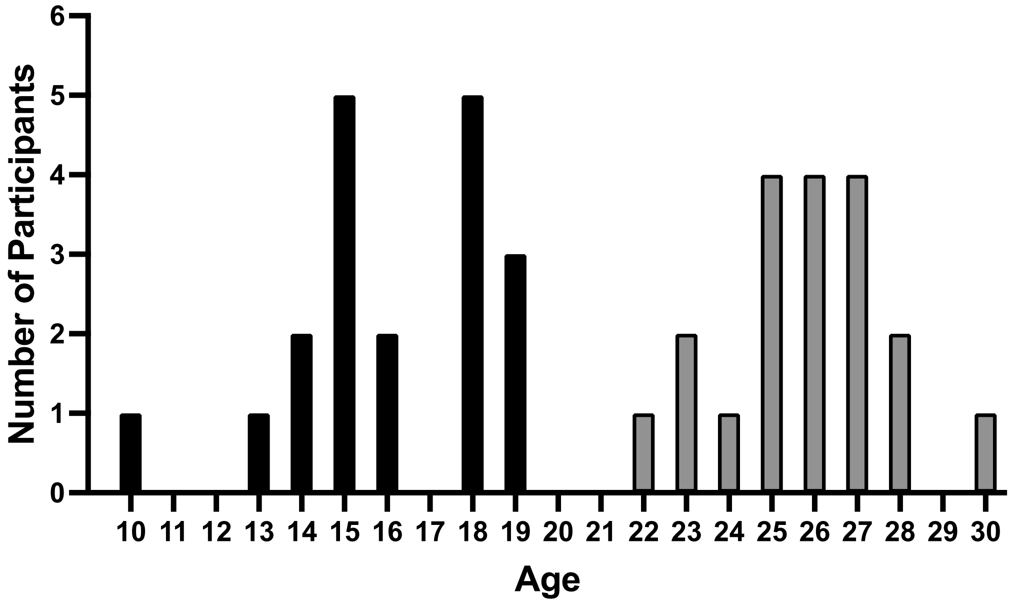


C

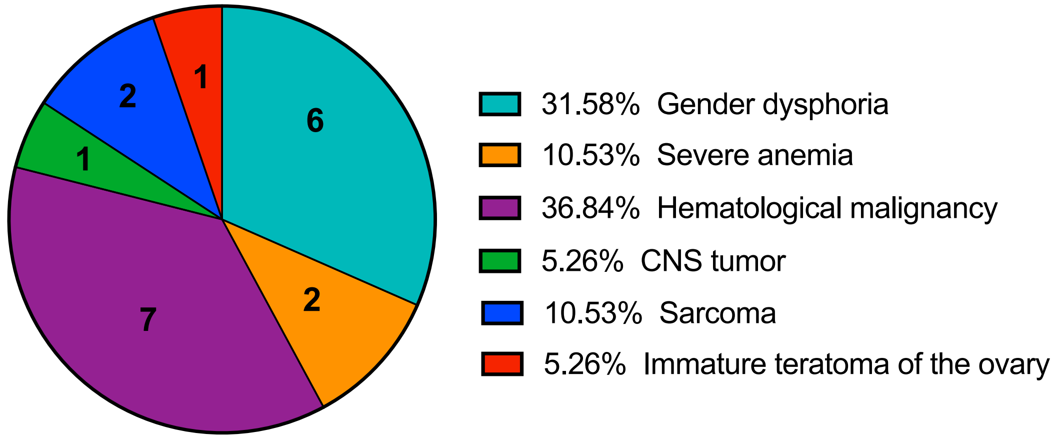


**Supplementary Figure 3. Demographics and IVF cycle characteristics (A), age: adolescents (black bars), oocyte donors (grey bars) (B) of participants, and medical diagnoses of adolescents (C) included in cumulus cell transcriptomic analysis.**

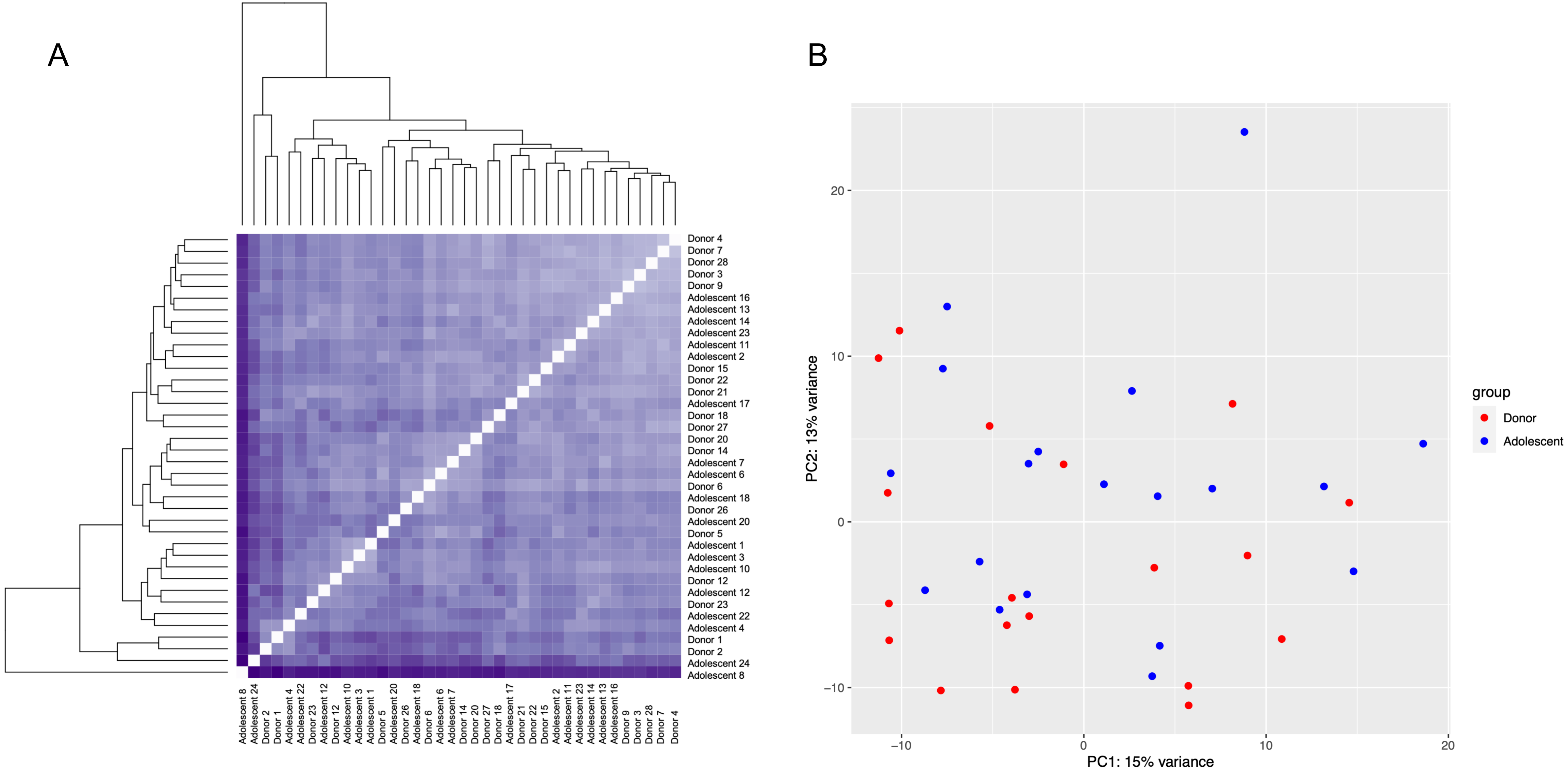


**Supplementary Figure 4. RNA-seq analysis of cumulus cells collected from adolescents and oocyte donors.**
A) Unsupervised hierarchical clustering B) Principal component analysis

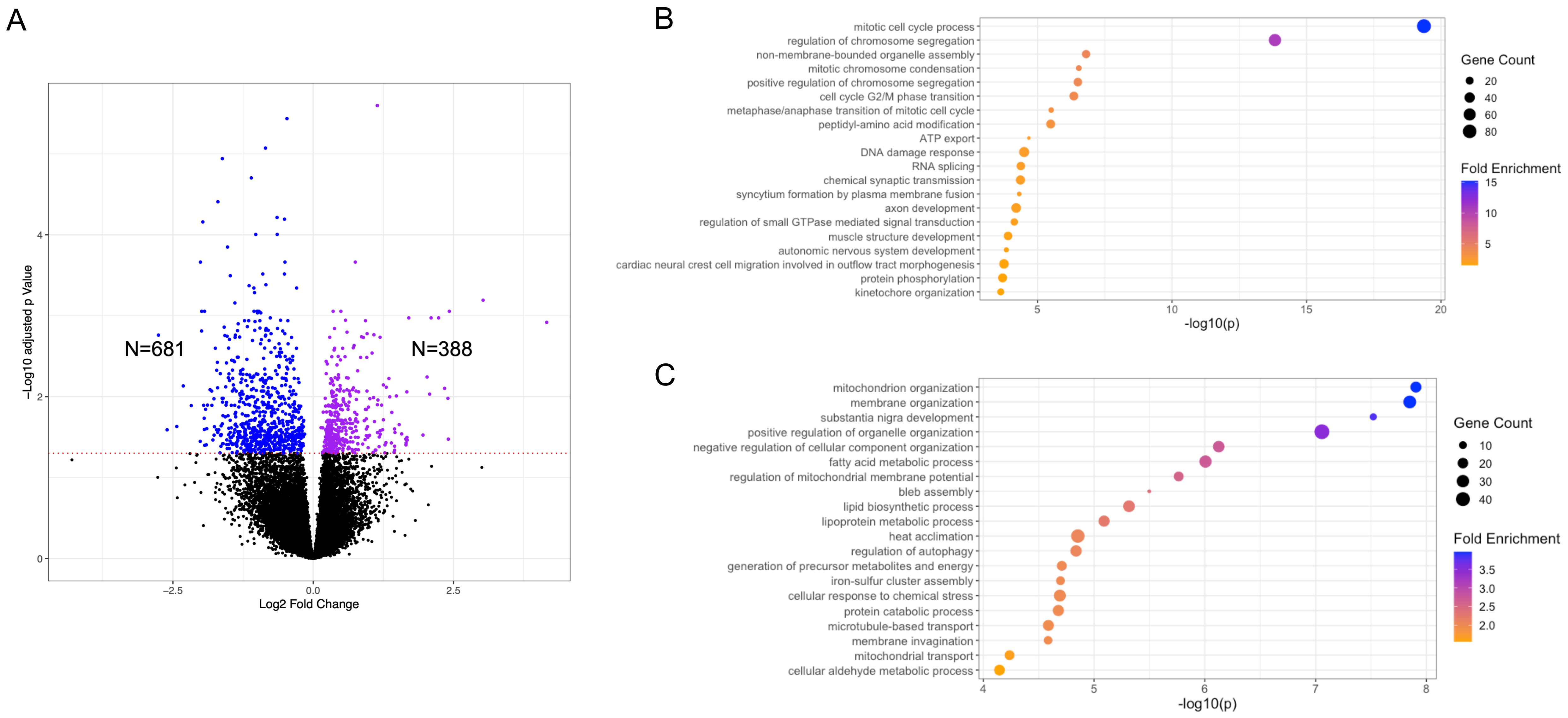


**Supplementary Figure 5. Comparative RNA-seq analysis of cumulus cells collected from adolescents <16 years old and oocyte donors.** A) Volcano plot shows downregulated (n=681) and upregulated DEGs (n=388) in adolescents (n=9) compared to donors (n=19) (dashed red line - adjusted p<0.05). B) Top 20 significantly downregulated GO terms in adolescents <16 years old compared to donors. C) Top 20 significantly upregulated GO terms in adolescents <16 years old compared to donors.

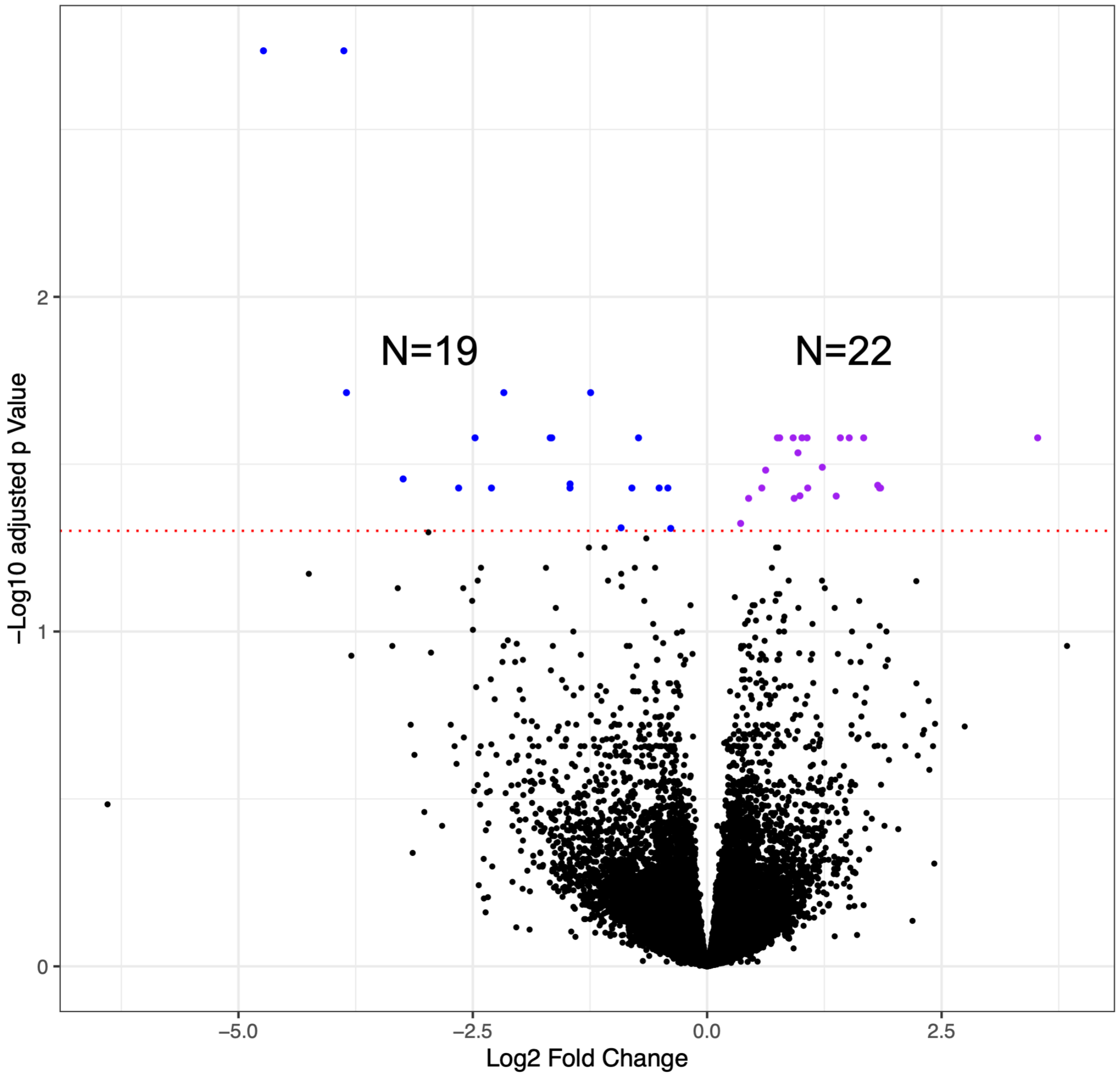


**Supplementary Figure 6. Comparative RNA-seq analysis of cumulus cells collected from adolescents ≥16 and <16 years old.** Volcano plot with downregulated (n=19) and upregulated DEGs (n=22) in adolescents ≥16 (n=9) compared to <16 years old (n=10) (dashed red line - adjusted p<0.05).

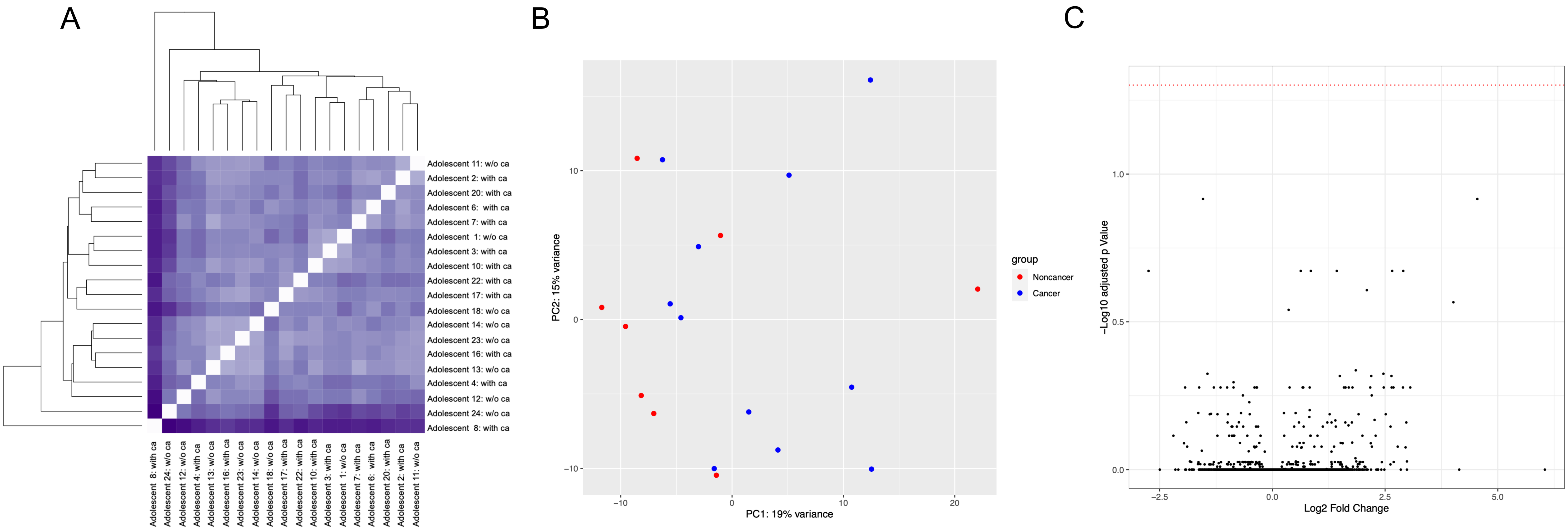


**Supplementary Figure 7. RNA-seq analysis of cumulus cells collected from adolescents with cancer compared to adolescents without the cancer diagnosis.** A) Unsupervised hierarchical clustering of adolescents with (n=11) and without (w/o) cancer (ca) (n=8) diagnosis B) Principal component analysis C) Volcano plot shows no DEGs between adolescents with and without cancer (dashed red line - adjusted p<0.05).

A

| | Adolescents (n=18) | Donors (n=16) | P value |
|---|---|---|---|
| **Age** (years) | 16.7 ± 0.6 | 27.3 ± 0.4 | <0.0001 |
| **BMI** (kg/m$^2$) | 25.1 ± 1.7 | 23.6 ± 0.5 | 0.4032 |
| **Race/ethnicity** (number of participants) | | | 0.2785 |
| Caucasian | 9 | 14 | |
| African American | 3 | 1 | |
| Asian | 1 | 1 | |
| Middle Eastern | 1 | 0 | |
| Caucasian-Hispanic | 2 | 0 | |
| **AMH** (ng/mL) | 3.2 ± 0.5 | 5.8 ± 0.6 | 0.0024 |
| **Antral follicle count (AFC)** | 16.5 ± 1.2 | 23.3 ± 1.6 | 0.0072 |
| **Luteal start** | 44.44% | 6.25% | 0.0189 |
| **Duration of stimulation** (days) | 11.3 ± 0.3 | 11.4 ± 0.3 | 0.8799 |
| **Number of monitoring visits** (days) | 6.1 ± 0.3 | 6.8 ± 0.3 | 0.1373 |
| **Type of USG** | | | <0.0001 |
| Transvaginal% | 38.89% | 100.00% | |
| Transabdominal% | 61.11% | | |
| **Total gonadotropin dose** (IU) | 5542 ± 471 | 4202 ± 435 | 0.0656 |
| **Peak estradiol** (pg/mL) | 2209 ± 234 | 3431 ± 303 | 0.0031 |
| **Number of oocytes** | 28.4 ± 3.4 | 29.9 ± 3.4 | 0.7506 |
| **Number of MIIs** | 19.4 ± 2.6 | 24.0 ± 2.9 | 0.1558 |
| **Number of MIs** | 1.6 ± 0.3 | 2.3 ± 0.4 | 0.2471 |
| **Number of GVs** | 3.3 ± 0.9 | 2.3 ± 0.6 | 0.4826 |
| **Number of degenerated oocytes at retrieval** | 1.5 ± 0.5 | 0.8 ± 0.3 | 0.1753 |
| **Number of EZs** | 2.1 ± 0.7 | 0.7 ± 0.4 | 0.0394 |

Values are presented as mean ± SEM. BMI = body mass index; AMH = Anti-Mullerian hormone; MIIs = mature metaphase II arrested oocytes; MIs = immature oocytes between GV and MII stage. GVs = immature oocytes with germinal vesicle; EZs = zona pellucida devoid of an oocyte.

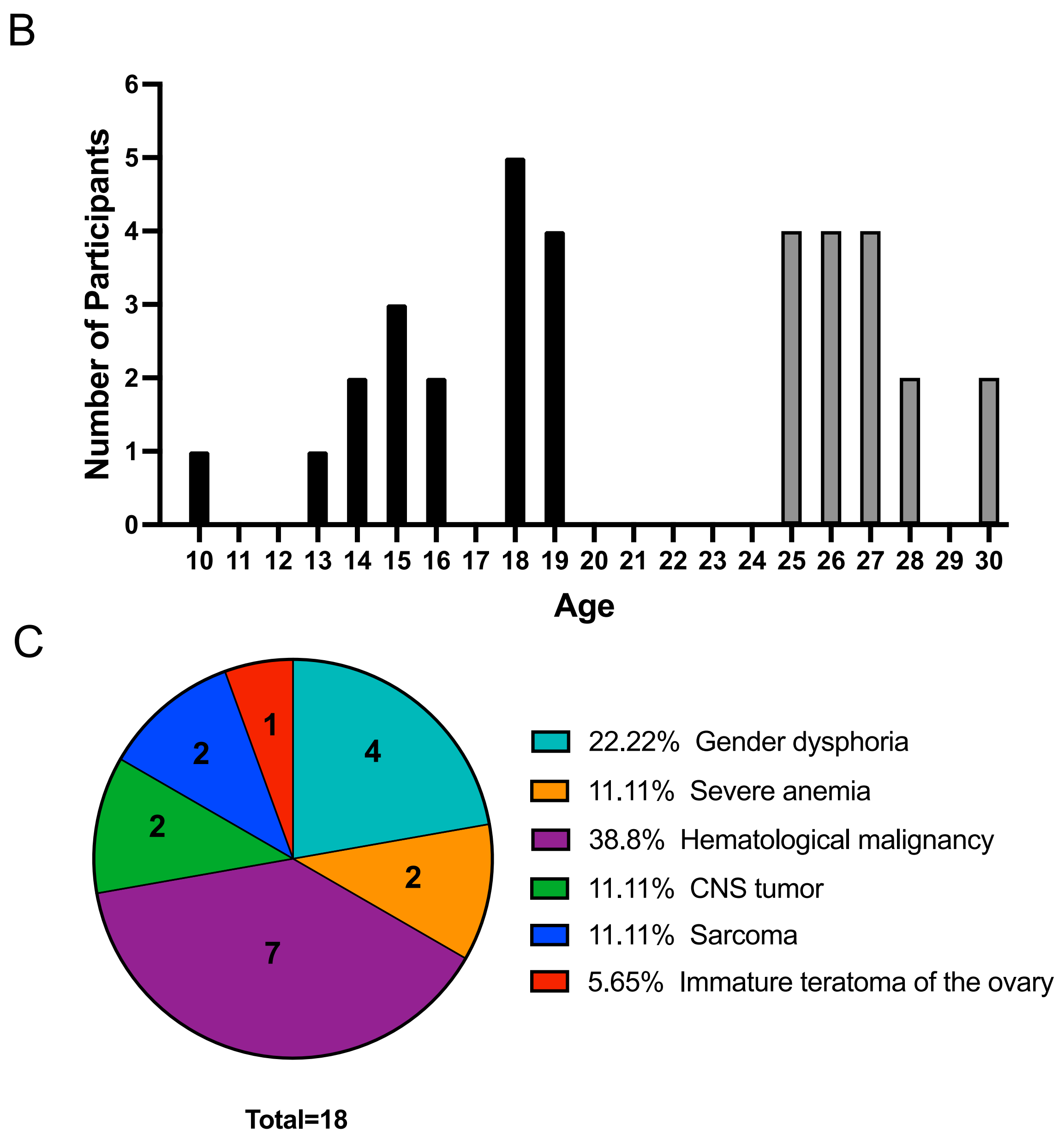


**Supplementary Figure 8. Demographics and IVF cycle characteristics (A), age: adolescents (black bars), oocyte donors (grey bars) (B) of participants, and medical diagnoses of adolescents (C) included in follicular fluid cytokine analysis.**

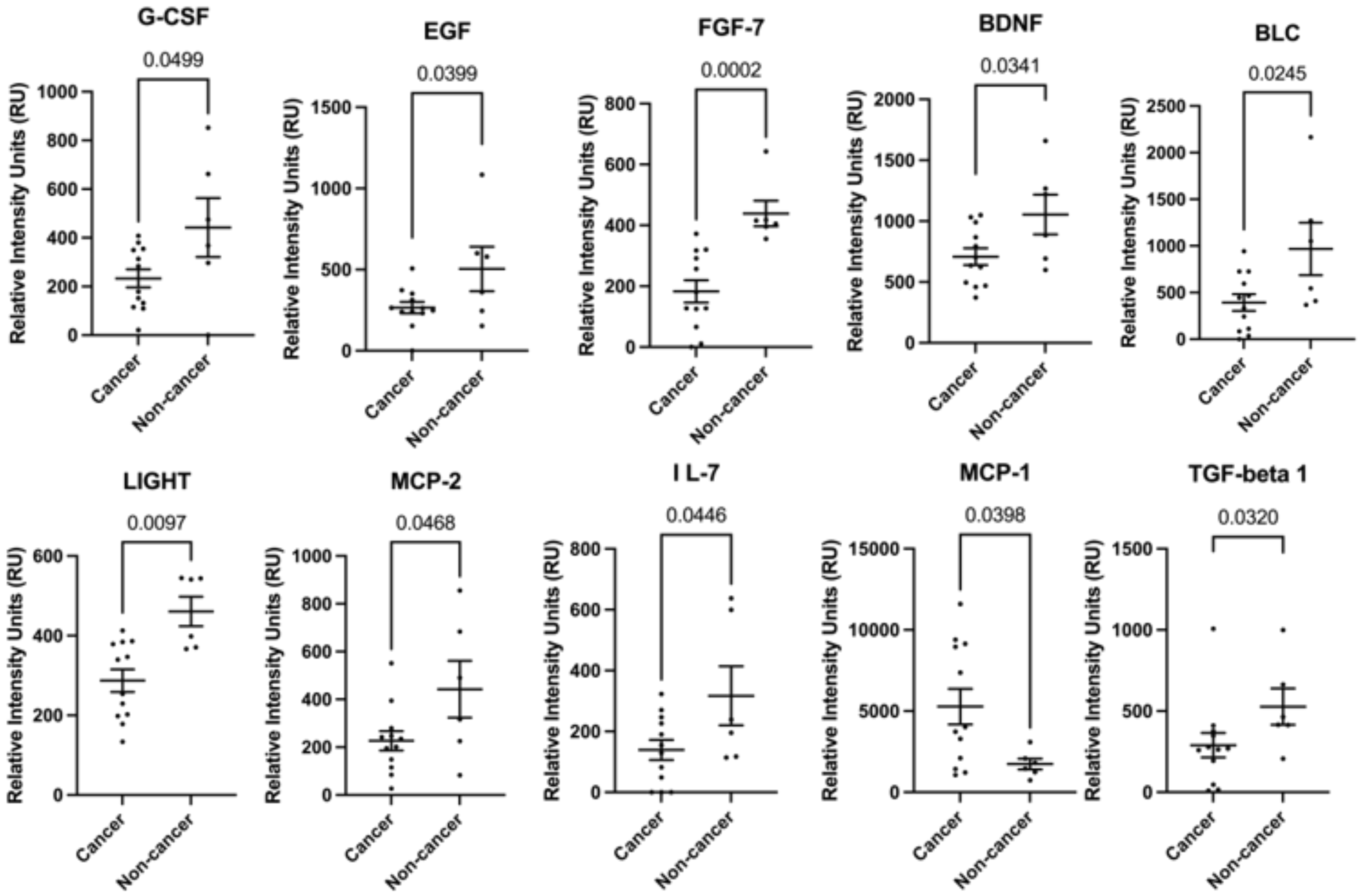


**Supplementary Figure 9. The comparison of follicular fluid cytokine levels in adolescents with and without cancer.** Analysis was performed to compare the levels of 80 cytokines in adolescent patients with cancer (n=12) compared to those without the cancer diagnoses (n=6). 10 out of 80 cytokines demonstrated significantly different follicular fluid levels.
